# The phylogenetic history of the *Gorteria diffusa* radiation sheds light on the origins of plant sexual deception

**DOI:** 10.1101/2022.12.22.521170

**Authors:** Boris Delahaie, Gregory Mellers, Roman T. Kellenberger, Mario Fernández-Mazuecos, Róisín Fattorini, Samuel F. Brockington, Allan G. Ellis, Beverley J. Glover

**Affiliations:** Department of Plant Sciences, University of Cambridge, Downing Street, Cambridge CB2 3EA, UK; CIRAD, UMR DIADE, F-34398 Montpellier, France; UMR DIADE, Université de Montpellier, CIRAD, IRD, Montpellier, France; Department of Biology (Botany), Faculty of Science, Universidad Autónoma de Madrid, Spain; Centro de Investigación en Biodiversidad y Cambio Global (CIBC-UAM), Universidad Autónoma de Madrid, Spain; Department of Biochemistry and Systems Biology, Institute of Systems, Molecular and Integrative Biology, University of Liverpool, Liverpool, L69 7ZB, United Kingdom; Department of Botany and Zoology, Stellenbosch University, Matieland, South Africa

## Abstract

The morphologically diverse daisy species *Gorteria diffusa* employs varying levels of sexually deceptive pollination. The species comprises at least fifteen spatially and phenotypically discrete floral morphotypes that are associated with a range of pollination strategies, from generalism to highly specialised sexual deception involving visual mimicry of females of the bee-fly *Megapalpus capensis*. However, the pattern of evolution of the unique floral traits in this lineage remains unknown because the phylogenetic history of the closely related floral morphotypes has proved unresolvable using traditional approaches. Here we apply genotyping-by-sequencing (GBS), a reduced representation sequencing technology that has significantly increased the tractability of phylogenetic problems involving recent radiations, to the recalcitrant phylogenetic problem of *Gorteria* across its South African distribution. Population genomic analyses show that individuals group according to morphotype, irrespective of geographic proximity, highlighting the distinctiveness of the morphotypes at the genetic level. We resolve the phylogenetic history of the closely related morphotypes, demonstrating that they are mostly well supported monophyletic entities that are grouped into at least three distinct geographically separated clades. Our results suggest that both incomplete lineage sorting and introgression across geographical clades have previously hindered reconstruction of the phylogeny of this species complex that has diversified rapidly during the Quaternary. Sexual deception is a phylogenetically derived pollination strategy within the complex that evolved at least twice, and was likely achieved by sequential evolution of a set of floral traits that in combination elicit sexual responses from the bee-fly pollinator. While insight into the evolution of sexual deception has been limited by strong phylogenetic conservatism of this strategy in other plant lineages, our results both provide the framework, and confirm the utility of *G. diffusa*, for further understanding the genetic pathways and selective pressures underlying the complex phenotypes required to exploit insect mating behaviour for pollination.

## Introduction

Evolutionary radiations are defined as rapid diversification events leading to the fast proliferation of the number of taxa in a clade. They are of particular interest to understand the variety of processes leading to adaptation and speciation, and have fascinated biologists for over a century. In order to decipher the processes involved in diversification and trace back the evolution of phenotypic diversity in radiations, reconstructing their evolutionary history is an important prerequisite. Recent evolutionary radiations are particularly interesting because the confounding effects generated by time in older clades (*e.g*. loss of the signal due to changes occurring after divergence) are not present (Whitfield and Lockhart, 2007). However, reconstructing their phylogenies is also a complicated task. When diversification occurs rapidly, lineages have limited time to accumulate substitutions that can be translated into phylogenetic signal (Shaffer and Thomson, 2007). Some regions of the genome remain undifferentiated, offering no phylogenetic resolution, while others present contrasting signals in the form of incongruent gene trees (*i.e*. showing different topologies). Two main biological processes are responsible for gene tree discordances: incomplete lineage sorting and introgression. First, when multiple speciation events occur close together in time, polymorphisms that are shared between the lineages might be retained by chance in some of these events but not others, therefore generating discrepancies between the gene trees and the species tree, a phenomenon called incomplete lineage sorting (hereafter ILS) (Degnan and Rosenberg, 2006). Second, it has recently become clear that introgression is ubiquitous across the tree of life (Mallet et al., 2016, *e.g*. Malinsky et al., 2018; Pease et al., 2018; Suvorov et al., 2021) and has various consequences for gene tree topologies. Generally, introgression between sister lineages will increase concordance between gene trees compared to an ILS-only scenario, but when introgression occurs between distant lineages, one type of discordant topology is expected to become more common (Hibbins and Hahn, 2021).

Fortunately, over the last decade high-throughput sequencing has enabled the simultaneous interrogation of large numbers of molecular markers. This data revolution has been accompanied by the development of various phylogenomic methods well suited for reconstruction and investigating the causes of gene tree discordance (see Bryant and Hahn, 2020; Hibbins and Hahn, 2021). While a number of problems remain to be solved (reviewed in Simion et al., 2020), these methods have proven useful in reconstructing the evolutionary history of various recent radiations that were previously considered as recalcitrant phylogenetic problems (*e.g*. Nater et al., 2015; Fernández-Mazuecos et al., 2018).

The Succulent Karoo biome, located in Southern Africa, is the only desert considered as a biodiversity hotspot, thanks to its exceptional floral richness and high rate of endemism (Cowling and Hilton-Taylor, 1994). It represents a recent component of the Greater Cape Floristic Region (Born et al., 2007), comprising lineages that have diversified in the last 10 my following Miocene aridification (Verboom et al., 2009). Interestingly, it hosts numerous taxa that are phenotypically and genetically diverse at the intra-specific level and for which taxonomic limits and evolutionary history remain poorly known, potentially highlighting a number of recent evolutionary radiations (*e.g*. Aizoaceae: Ellis et al., 2007, 2006; Klak et al., 2003; Musker et al., 2021). Among them, the daisy *Gorteria diffusa* Thunb. stands out as an extraordinarily diverse species complex, presenting at least 15 distinct morphological forms (morphotypes) distributed in parapatry (Fig. 1). These morphotypes vary in flower colour as well as in the presence and complexity of anthocyanic spots on their ray florets (Fig. 1) (Ellis et al., 2014). Another striking characteristic of the species is that it is considered one of a very few sexually-deceptive plant species that are not orchids (Johnson and Midgley, 1997; Ellis and Johnson, 2010). The males of its main pollinator species, the bee fly *Megapalpus capensis* Wiedemann, are tricked by the resemblance of the flower spots to congeners so that they search for and attempt to copulate with these imitation females. Variation in the extent of sexual response to different morphotypes has been shown to be tightly linked to the differential spot morphologies (Ellis et al., 2014; Ellis and Johnson, 2010), and varies from no sexual behaviour at all, *i.e*. only feeding, to pseudo-copulation with the flower. The floral morphologies which appear most successful in mimicking female pollinators comprise strongly integrated trait combinations, including UV reflective highlights and raised papillate sections of epidermis within the spot (de Jager and Ellis, 2012; Ellis et al., 2014). Higher levels of pollen export have been shown to occur as the extent of sexual deception increases, suggesting that this pollination strategy has a significant role in plant fitness and is potentially a key factor in the reproductive isolation of morphotypes and the recent diversification of the species (Ellis and Johnson, 2010).

**Figure 1:**
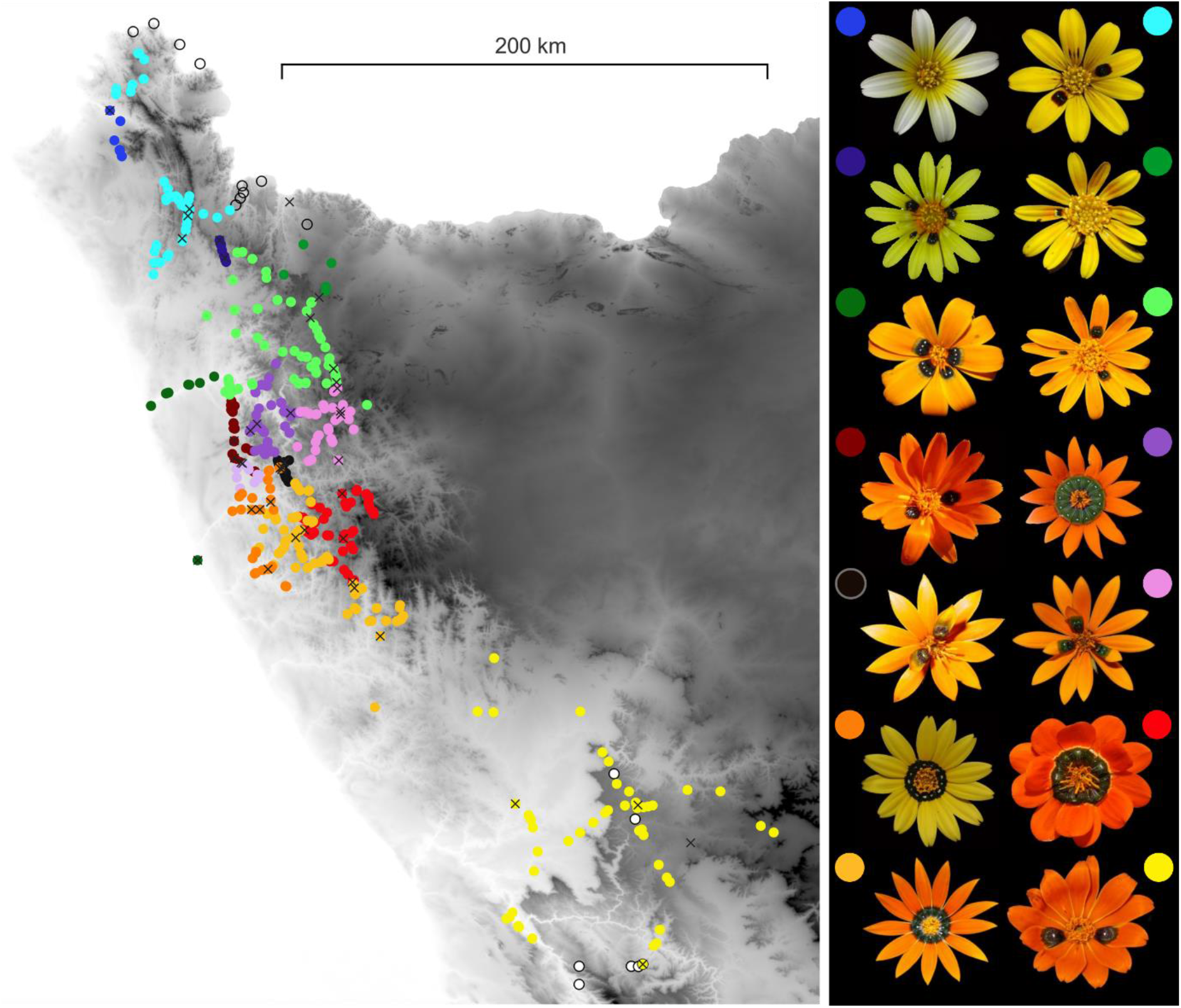
Distribution range of the 15 different morphotypes of *Gorteria diffusa* across Namaqualand, South Africa. Each dot represents a sampling location where morphotypes have been determined. Crosses represent sampling localities for this study. The standard floral phenotypes of the 14 morphotypes included in this study are illustrated. Coloured dots alongside the pictures link to the map colours. Morphotype names are as follows, clockwise from the top left: Khubus, Rich, Stein, Okiep, Naries, Spring, Cal, Nieuw, Garies, Soeb, Buffels, Koma, Kleinzee (Kz), Roub. Localities of *Gorteria corymbosa* (open circles) and *G. personata* (white dots) are indicated in areas where their ranges abut/overlap with *G.diffusa*.

To explore the evolution of this intriguing and highly successful strategy, and to understand the processes that have led to these floral diversification events, reconstructing the evolutionary history of this species complex is necessary. However, previous attempts at reconstructing the evolutionary history of *G. diffusa* (Stångberg et al., 2013a) have been unsuccessful, suggesting that its diversification is too recent to be resolved with classic phylogenetic markers. In this study, we used a high-throughput sequencing technique (Genotyping-by-Sequencing, hereafter GBS; Elshire et al., 2011) to recover thousands of markers with the aim of delineating the genetic boundaries between the morphotypes, understanding their evolutionary relationships and reconstructing patterns of floral morphological evolution in this species complex. Specifically, we combined population genetic tools, concatenation and coalescent-based phylogenetic reconstruction methods to understand its evolutionary history. Additionally, we tested for gene flow between the different morphotypes and performed ancestral trait reconstruction to shed light on the origin of sexual deception in the species. We obtain a well-resolved phylogeny and conclude that sexual deception arose twice within the species complex, most likely as a result of combining traits already present in distinct genotypes. Building on these results, we also highlight some perspectives regarding the role of hybridization and the evolution of reproductive isolation in this system.

## Material and Methods

### Sampling scheme

We sampled a total of 96 plants from the 14 morphotypes of *G. diffusa* as described in (Ellis and Johnson, 2009; Stångberg et al., 2013a), and one additional recently discovered morphotype (Roub; Ellis (unpublished)). In order to tease apart the relative contributions of geography and phenotypic characteristics to the genetic structure of the species, two to three localities of each morphotype were sampled, spanning a representative proportion of their range (Fig. 1). Additionally, we sampled other closely related *Gorteria* species (*G. piloselloides* (Cass.) Stångb. & Anderb., *G. corymbosa* DC., *G. alienata* (Thunb.) Stångb. & Anderb. and *G. personata* L.) and one individual from the sister genus *Hirpicium (H. echinus* Less.) to serve as outgroup taxa and to confirm previous findings regarding their relationships to *G. diffusa* (Stångberg et al., 2013b). For each individual, a few leaves were collected and subsequently dried in silica gel. Correct identification was ensured by sampling individuals in full flower so that the phenotypic characteristics could be checked and compared to the morphotype descriptions (found in Ellis and Johnson, 2009b; Stångberg et al., 2013a). For each population sampled, herbarium replicates are available from the University of Cambridge or Stellenbosch University Herbaria (barcodes CGE14001-CGE14122). All sampling was conducted under research and collecting permits issued by Northern Cape Department of Environment and Nature Conservation and Cape Nature.

### DNA extraction and genotyping-by-sequencing

For all individuals, genomic DNA was extracted using a modified version of the CTAB protocol (Doyle and Doyle, 1987, see supp. mat. for details). Genotyping-by-Sequencing library preparation was done in-house following Fernández-Mazuecos et al., 2018 (originally adapted from Escudero et al. (2014), see supp. mat. for protocol details). We chose the restriction enzyme *PstI* as it is a relatively rare cutter (CTGCA|G) allowing us to target a sufficient sequencing depth considering the relatively large genome size of *G. diffusa* (ca. 2Gb estimated through flow cytometry, unpublished data). This enzyme is sensitive to methylation and therefore tends to target hypomethylated regions, avoiding the repetitive fraction of the genome (Kilian et al., 2012). Barcodes were designed to avoid potential misidentification and to ensure nucleotide diversity and balance for sequencing. The sequencing was performed on a single lane of 100bp paired-end on an Illumina HiSeq 2000 by the Beijing Genomics Institute (Copenhagen, Denmark).

### Assembly and filtering of the data

The quality and presence of Illumina adapters were assessed using *multiQC* (version 1.9, (Ewels et al., 2016)). *Trim_galore* (version 2.1 with *Python* 3.7.3 (Martin, 2011)) was used in order to remove adapters and low quality fragments and to trim the reads to 90bp. Cleaned and trimmed reads were then demultiplexed using the function *process_radtags* from *Stacks* (version 2.41 (Rochette et al., 2019)). We used a good quality unpublished draft assembly of the genome of *G. diffusa* (Spring morphotype, 4222 contigs with an NG50 length of 0.43 Mbp, a total haploid size of 0.97 Gbp and 95.7% estimated gene completeness) to aid the assembly of the GBS data. *BWA-MEM* (version 0.7.17-r1188, Li and Durbin, 2009) was used to map individual sequences against this reference genome and to produce SAM files using default options. *samtools* (Li et al., 2009) was used to sort and compress the SAM files into BAM ones. SNPs were then called using the wrapper script (*ref_map.pl*) of the Stacks program (version 2.41, Rochette et al., 2019).

Filtering parameters, such as the amount of missing data tolerated in a dataset, have been shown to influence phylogenetic reconstructions (*e.g*. Leaché et al., 2015; Lee et al., 2018; Wiens and Morrill, 2011). For this reason, all inferences were repeated on various datasets filtered for different amount of missing data (minimum individual coverage of the loci, varying from 10% to 90% with a 10% increment). In each dataset, we used *VCFtools* (version 0.1.15, Danecek et al., 2011) to filter out loci having an average read depth lower than 8 across all genotypes in order to further reduce the presence of erroneous genotype calls.

### Population structure

In order to characterize population structure at the intraspecific level, we used two complementary methods to visualize genetic structure associated with geographic and morphotype differentiation. To do so, we used a reduced dataset where loci genotyped in less than 80% of the individuals were removed. We also applied a minor allele count threshold of 5 to ensure rare alleles were filtered out. We then used the R package *SNPRelate* (Zheng et al., 2012) to compute Identity-By-State (IBS) estimates in order to remove potentially highly related individuals that could affect our analyses. For any pair of individuals with an IBS value greater than 95%, the individual with the highest amount of missing data was excluded.

First, we performed a PCA using the *R* package *adegenet* (Jombart and Ahmed, 2011) after having thinned the reduced dataset to remove markers in linkage disequilibrium using the software *plink* (with the parameters --indep 50 5 2; Purcell et al., 2007). Second, we used the program *fineRADstructure* (Malinsky et al., 2018b) to infer population structure based on shared ancestry among all individuals. As co-ancestry is based on whole haplotypes, we did not remove SNPs in linkage disequilibrium. We used a total of 200,000 MCMC iterations with a burn-in period of 100,000.

### Phylogenetic reconstruction

In order to obtain a robust phylogenetic hypothesis for the relationships within *G. diffusa*, we used two complementary phylogenetic reconstruction methods based on concatenation and coalescence. Since recent gene flow can have a strong influence on the reconstruction of phylogenetic trees (Eaton et al., 2015; Leaché et al., 2014; Nadeau et al., 2014), we removed 8 individuals sampled from 4 localities that were very close to contact zones (< 2 km) and likely influenced by contemporary gene flow (see Fig. 1) as well as 3 individuals inferred as highly related (see previous section).

### Concatenation approach

We performed maximum-likelihood phylogenetic tree estimations using *iqtree* (version 2.0.3, (Minh et al., 2020)). For each missing data threshold, we built concatenated matrices of the whole loci (SNPs and their flanking regions, as recommended in (Leaché et al., 2015)). The best models of nucleotide substitution were selected using ModelFinder (Kalyaanamoorthy et al., 2017) within iqtree based on Bayesian Information Criterion (BIC). To assess branch support, we computed both ultrafast bootstrap (UFB, Hoang et al., 2018; Minh et al., 2013) with 1000 replicates and an approximate likelihood ratio test with the nonparametric Shimodaira-Hasewaga correction SH-aLRT with 1000 replicates (Guindon et al., 2010). To interpret these results, we used a conservative approach to characterize strong support for any given node by using a 95% threshold for UFB and a 80% threshold for SH-aLRT (following recommendations in (Minh et al., 2013)).

### Coalescent-based approach

Since the diversification of *G. diffusa* is thought to be recent (Stångberg et al., 2013a)), we used a coalescent-based approach relying on quartet trees, ASTRAL-III (Zhang et al., 2018), to further assess the relationship between the morphotypes. This method is known to better incorporate incomplete lineage sorting than the concatenation-based approach and relies on the assemblage of gene trees into species trees, therefore enabling the study of the concordance in the phylogenetic history observed in different genes. Although ASTRAL-III performs best with longer fragments, it has been shown to be reliable with short fragments as well (Chou et al., 2015). First, we extracted individual alignments for each locus using a custom script based on the *populations* function of Stacks. Second, in order to reconstruct individual gene trees, we used the ParGenes tool (Morel et al., 2019) enabling highly parallelized gene tree inference using RAxML-ng (Kozlov et al., 2019). For each gene tree, we performed 20 tree searches using 10 random and 10 parsimony-based starting trees. The GTR+F+G4 substitution model was chosen and we performed 100 parametric bootstraps for each gene tree to assess support. Then, as recommended by (Zhang et al., 2017), we collapsed nodes with very low support (non-parametric bootstrap < 10%) in each gene tree using Newick utilities (Junier and Zdobnov, 2010) before building the final trees. For each missing data threshold, we built a tree using the multi-individual mode (Rabiee et al., 2019) where individuals are grouped *a priori* into putative species (here corresponding to recognized species or *G. diffusa* morphotypes). Using multiple individuals per species and defining species boundaries *a priori* is beneficial to retrieve the correct species tree (Rabiee et al., 2019). However, when there are uncertainties in the *a priori* species delimitation, the inferred species tree may not be correct. To overcome this possibility, as recommended by Rabiee et al. (2009), we repeated this analysis using sampling sites as the *a priori* grouping to validate the monophyly of the different species/morphotypes.

All models were run using default parameters. In order to reliably characterize the support for each node, we computed ASTRAL support values, which can be interpreted as local posterior probabilities (Sayyari and Mirarab, 2016).

### Tree comparisons

We compared both the support and the topology of the different trees obtained. First, regarding support, we simply compared the distribution of the support values obtained for each tree. Second, we compared the different topologies obtained for the different analyses and datasets using the clustering information distance described in (Smith, 2020a). These metrics were estimated using the R-package TreeDist (Smith, 2020b) and a multidimensional scaling approach was used to visualize distances between the different trees using the ‘cmdscale’ function in R.

### Investigating causes of gene tree discordance

We used D-suite (Malinsky et al., 2021) to formally test for introgression between clades by estimating the D statistic (also known as the ABBA-BABA statistic) (Durand et al., 2011). In a quartet tree topology (((P1, P2), P3), O), O is the outgroup and is used to define the ancestral allele called A, while the derived alleles are noted B. Three different topologies can be observed. BBAA corresponds to a situation with complete lineage sorting without introgression where P1 and P2 share the derived allele (B). In a situation with incomplete lineage sorting and/or introgression, ABBA and BABA topologies can be observed and correspond respectively to P2 and P3 sharing the derived allele, and P1 and P3 sharing the derived allele. Under a model without introgression, ABBA and BABA topologies are expected to show equal frequencies. In an introgression scenario, ABBA is expected to show higher frequency than BABA if there is introgression between P2 and P3, while BABA is expected to show higher frequency than ABBA if there is introgression between P1 and P3. The D statistic corresponds to the normalized difference between the number of ABBA and BABA patterns. D is therefore bounded between −1 and 1 and is expected to equal 0 when there is no introgression.

To do so, we used the same dataset as in the phylogenetic reconstruction analyses and applied a minor allele count threshold of 5 to ensure rare alleles were filtered out. Since D-suite can handle missing data, we used the most complete dataset. D-suite is implemented to work directly on VCF files so we used the *populations* function of *Stacks* to extract it. Our focus was on introgression between morphotypes of *G. diffusa* but we also explored introgression between *G. diffusa* and its two closely related sister taxa, *G. personata* and *G. corymbosa*, by including them in the analysis. We used *G. piloselloides* as the outgroup. Each morphotype was considered as a distinct species in the analysis. We used the function Dtrios from D-Suite (Malinsky et al., 2021) which estimates D statistics for all combinations of trios. To test whether D significantly differed from 0, Z-scores and associated *p*-values were estimated. As each species/morphotype is involved in multiple tests, we then corrected the *p*-value using the Holm-Bonferroni method to account for multiple comparisons.

### Ancestral state reconstruction

In order to evaluate the evolution of sexual deception in *Gorteria diffusa*, we used three discrete floral traits that represent the bulk of floral variation in the system and are known to be responsible for variation in pollinator behaviour: ray floret colour (coded as yellow or orange), number of anthocyanin spots (coded as absent, few -when present only on 1 to 5 ray florets- or full ring -when present on all ray florets) and the presence of anthocyanin spot papillae (coded as present or absent). Additionally, we directly mapped the behaviour of the main pollinator on the different morphotypes according to three different categories: feeding, inspection and pseudo-copulation. These traits were obtained from previous work on the species (Ellis et al., 2014; Ellis and Johnson, 2009). We performed a stochastic character mapping analysis (Huelsenbeck et al., 2003) using the R-package PHYTOOLS (Revell, 2012). We used a tree built on a reduced dataset containing only one individual per morphotype. In order to obtain the best topology possible, we chose the individual with the highest number of loci retrieved in each morphotype. This tree was built using the same concatenation approach as for the full dataset. We chose to use the concatenated tree as branch length is expected to be more robust than for the coalescent-based approach. For each trait with two states, we tested two models of trait evolution: (i) ER, where rates are equal for all transitions among states, and (ii) ARD, where all rates are different. For traits with three states, we tested three models of trait evolution: (i) ER, (ii) ARD and (iii) SYM, where transition rates vary but backward and forward rates for each transition are equal. For each trait, we selected the best model using the Akaike Information Criterion (AIC). We then based our reconstruction on 1,000 stochastic reconstructions.

### Phylogenetic dating

To estimate a temporal framework for the divergence of *G. diffusa*, nuclear ribosomal internal transcribed spacer (ITS) and plastid *trn*L-*trn*F sequences of 12 morphotypes of *G. diffusa* and 152 additional species of Cichorioideae representing all major lineages were obtained from the GenBank database (Funk and Chan, 2008; Howis et al., 2009; Stångberg et al., 2013b; among others; Table S2). Sequences of two additional morphotypes of *G. diffusa* (Spring and Stein) were newly generated. Alignments were conducted using MAFFT 7 (Katoh and Toh, 2008), and substitution models were determined under the Akaike Information Criterion in jModelTest 2.1 (Darriba et al., 2012). Divergence times were estimated based on a concatenated matrix using the relaxed molecular clock method implemented in BEAST2 (Bouckaert et al., 2014). Fossil calibrations within the Cichorieae were implemented following (Tremetsberger et al., 2013). Specifically, the minimum ages of *Cichorium-type* (22 Ma) and *Sonchus-type* (5.4 Ma) fossil pollen (Blackmore et al., 1986; Hochuli, 1978) were set as minimum stem age constraints for clades where these pollen types are found (see Tremetsberger et al., 2013 for details). Additionally, a secondary calibration was set for the root node (uniform prior with bounds 26-35 Ma) following the estimate of the TimeTree database (Kumar et al., 2017), which is based on multiple previously-published studies. A birth-death model was set as tree prior, and substitution rate variation was modelled using a lognormal distribution. Four Markov chain Monte Carlo analyses were run for 100 million generations each, with a sample frequency of 10000. Parameter analysis in Tracer showed adequate chain length. Chains were combined using LogCombiner, after discarding the burn-in, and trees were summarized in a maximum clade credibility (MCC) tree obtained in TreeAnnotator and visualized in FigTree.

## Results

### Genotyping-by-sequencing and data assembly

The genotyping-by-sequencing experiment yielded 244 million clean reads. 12 individuals were removed beforehand due to very low coverage (<200,000 reads) and 4 due to problems with barcodes. The remaining 80 individuals had an average of 3 million clean reads per individual, with a minimum of 311,000 and a maximum of 9.1 million reads. Individuals with low number of reads or sequencing depth did not belong to any particular population or morphotype, and were likely the result of low DNA quality or technical problems during multiplexing. The number of loci retained was highly variable depending on the missing data proportion tolerated, varying from 491 with 10% to 4419 for the most complete dataset (details can be found in Table 1).

**Table 1:** Number of loci reconstructed for each dataset.

| Missingness | Number of loci |
| --- | --- |
| 10 | 491 |
| 20 | 852 |
| 30 | 1178 |
| 40 | 1497 |
| 50 | 1817 |
| 60 | 2212 |
| 70 | 2653 |
| 80 | 3261 |
| 90 | 4419 |

### Population structure

The PCA performed on the reduced dataset revealed a clear genetic structure at the within-species level and supported the premise that the morphotypes represent distinct genetic entities (Fig. 2). The different individuals sampled in different sites from the same morphotype tended to cluster together. At one end of the first axis, Rich and Khubus, the northernmost morphotypes, were isolated from the rest of the morphotypes. At the other end of the first axis, 9 morphotypes were separated in an approximate North-South gradient along the second axis. The morphotypes Okiep, Stein, Kleinsee and Roub were found in between these two groups. Interestingly, although the different morphotypes were globally structured according to geography in a North-South gradient, some groupings were at odds with geography, further highlighting the distinctiveness of the morphotypes at the genetic level. The individual sampled from the Kleinsee morphotype, for example, was associated with the morphotypes Okiep and Stein on all principal components. However, Kleinsee is in much closer geographical proximity to Soeb and is located more than 100km away from Okiep and Stein (Fig. 1, Fig. 2a, Fig. S1). Similarly, although being sampled at least 190km from any other morphotype, the morphotype Nieuw clustered with the other morphotypes from the South of the distribution on the first four PCs. Some individuals sampled close to hybrid zones also showed interesting patterns, as they tended to cluster with their respective morphotypes irrespective of geographic proximity. This was the case for Spring 1 and Okiep 1 (ca. 3 km apart), Cal 2 and Garies 1 (ca. 3km apart) and Buffels 3 and Soeb 1 (ca. 1km apart) (Fig. 1, Fig. 2a, Fig. S1).

**Figure 2:**
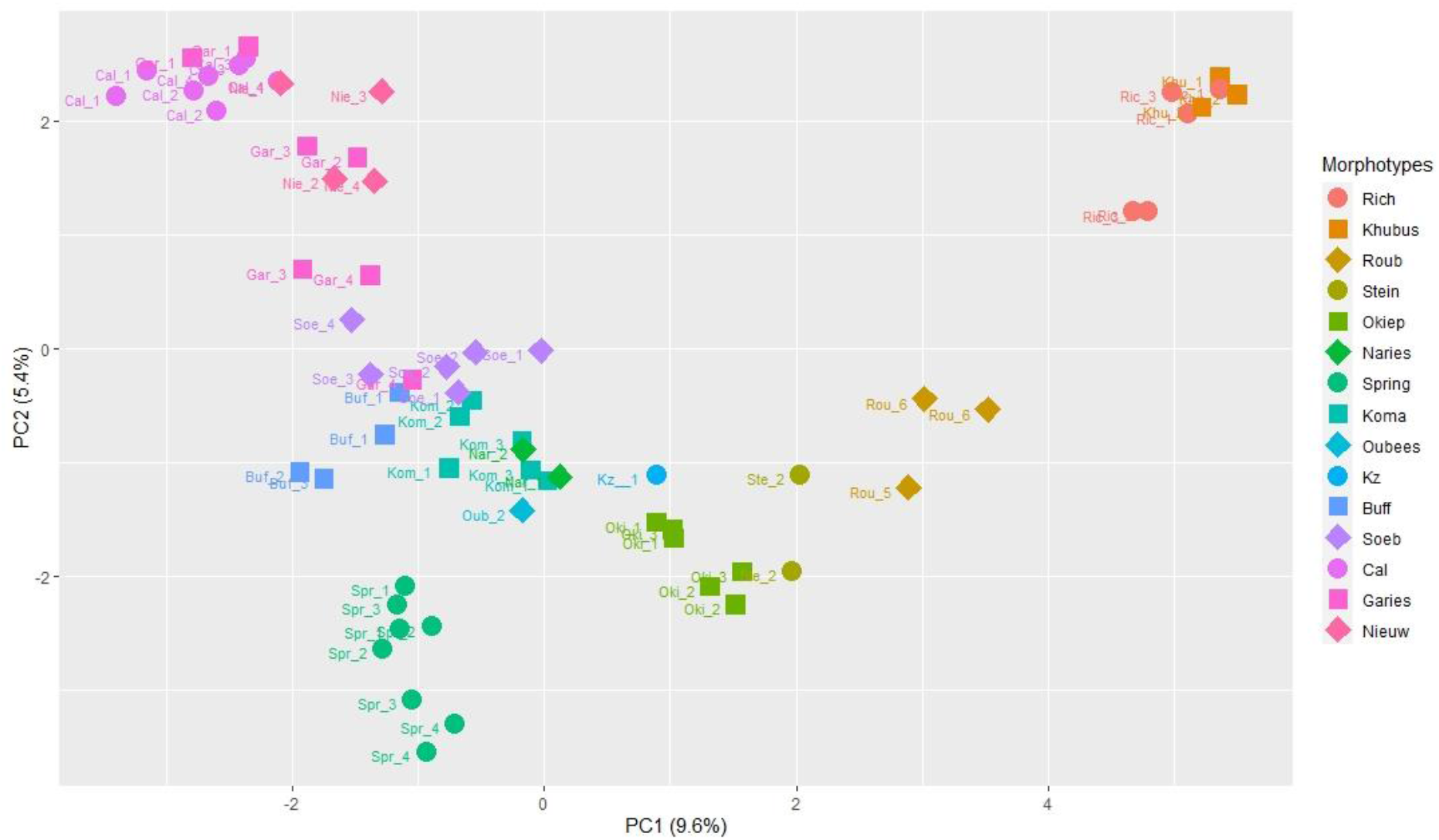
Principal components analysis of the 824 SNPs shared by 80% of the *Gorteria diffusa* individuals. Outgroup species were not included in this analysis. Colour and shape correspond to the different morphotypes, and the name of each sampling locality is specified next to the position of each individual (shortened to the 3 first letters of the morphotype). First and second axis are represented, corresponding respectively to 9.6% and 5.4% of explained variance.

The *fineRADstructure* analysis based on co-ancestry highlighted similar patterns to the PCA with each morphotype forming clusters of high co-ancestry (Fig. S1). The three main morphotype groups highlighted in the PCA were also apparent in this analysis. Interestingly, the three individuals from the morphotype Roub showed affinities with two groups: Rich-Khubus and Okiep-Stein-Kleinsee. Roub is phenotypically closer to the Okiep-Stein-Kleinsee morphotypes but is found geographically closer to the range of the Northern morphotypes (ca. 15k from Rich), which could be an indication of hybrid ancestry between these two groups.

### Phylogenetic reconstruction

#### Concatenation-based approach

Of the 9 topologies obtained using the concatenation approach, the ones built with the highest number of loci consistently showed the highest support values (Fig. S2A and S2B). Support tended to decrease for missing data levels lower than 70%. Hereafter, we focus on results using the datasets with highest support values (from 90 to 70% missing data).

First, all topologies confirm the monophyly of *Gorteria diffusa* with *G. corymbosa*, *G. personata*, *G. piloselloides*, *G. alienata*, and *Hirpicium echinus* being well supported as distinct from *G. diffusa*. This also confirms the results of (Stångberg et al., 2013b) that *G. piloselloides*, which was until recently considered as a morphotype of *G. diffusa* (called ‘Worcester’ in Ellis & Johnson, 2009), is a distinct lineage. *Gorteria corymbosa* appears as sister to the *G. diffusa* radiation, a relationship that was uncertain in previous work where either *G. personata* or *G. corymbosa* were inferred as the closest relative to *G. diffusa*.

Second, these trees were well supported overall at the within *G. diffusa* level and reveal strong phylogenetic structure between the different morphotypes of *G. diffusa* with all morphotypes inferred as monophyletic in most datasets, suggesting that they represent distinct groups. However, two out of the 15 morphotypes included in this study (Garies and Naries) were inferred as polyphyletic in some datasets. Interestingly, the pairs of morphotypes that are known to hybridize: Cal-Garies, Garies-Soeb, Soeb-Buffels, Koma-Soeb, Spring-Okiep, Spring-Naries (Fig. 1) were all inferred in distinct clades and were not necessarily found closely related in the topology, further confirming that they form clear and distinct entities. In the well-supported topologies (between 70% and 90% missing data), a single morphotype-level topology was found (Fig. 3a, Fig S3). As the level of missingness decreased, most of the among-morphotype relationships were largely congruent and only the positions of Spring, Buffels, Naries and Koma were variable. Interestingly, the nodes involving these morphotypes were generally the least supported ones in the best-supported trees, further highlighting the uncertainties surrounding their phylogenetic position.

**Figure 3:**
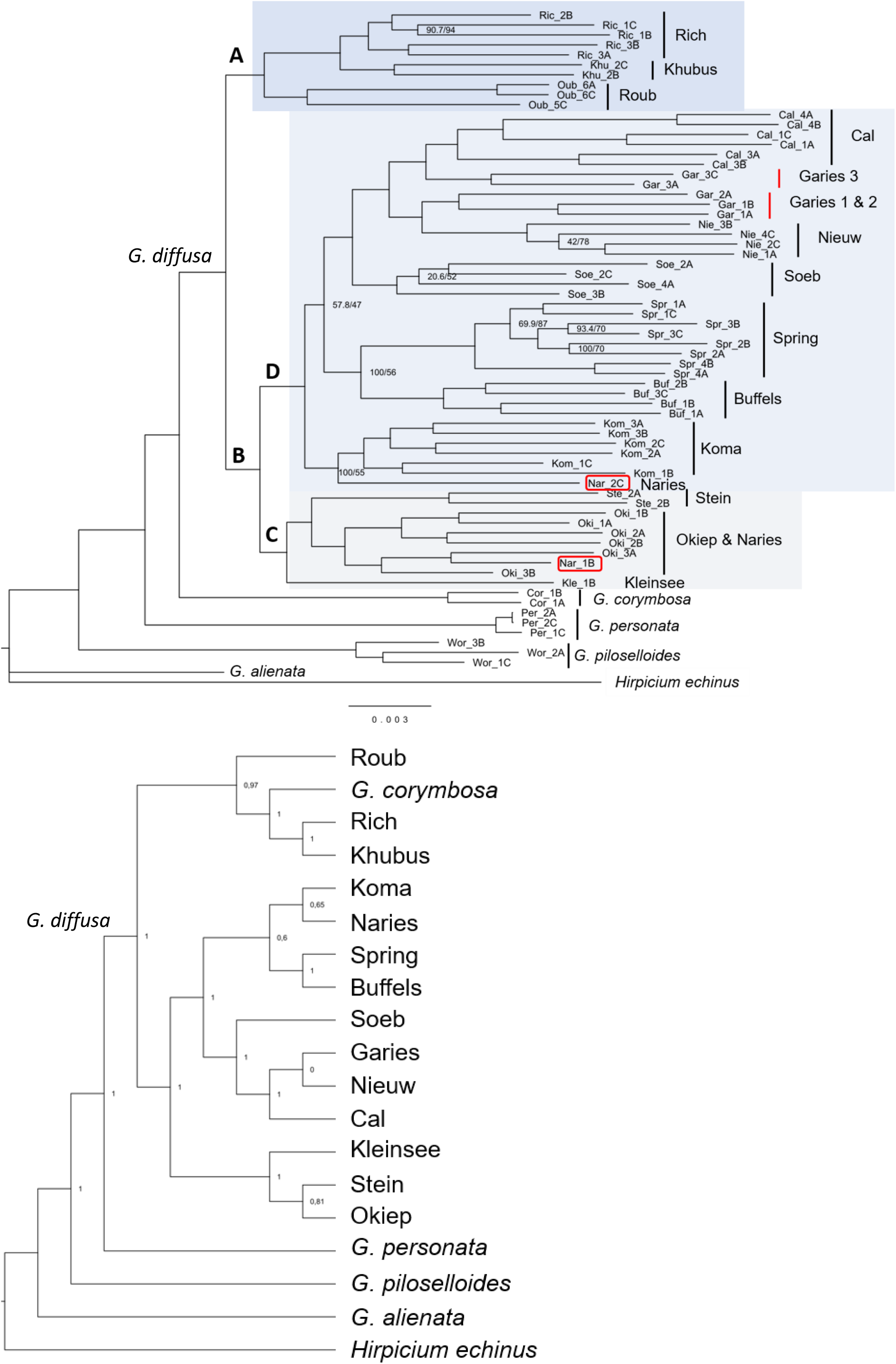
**a.** Phylogeny obtained with the concatenation-based approach on the 90% missing data dataset. Support values at each node correspond to ultra-fast bootstrap / SH-aLRT based on 1000 replicates each. Support values are not reported for well-supported nodes (UFB > 95, SH-aLRT >= 80). Clades mentioned in the main text are depicted as large letters and highlighted with various shadings. **b.** Phylogeny obtained with the coalescent-based approach on the 90% missing data dataset. Support values at each node correspond to local posterior probabilities as computed by the Astral-III method. In this analysis, individuals are grouped in putative species *i.e*. the recognized outgroup species and the different morphotypes.

The most basal split within *G. diffusa* involved a Northern clade (clade A, Fig. 3a) comprising the morphotypes Rich, Khubus and Roub occurring in the Richtersveld area (Fig. 1) and a clade (‘B’) containing the diversity found in the rest of Namaqualand. Within clade B, the first split involved a clade (‘C’) containing four morphotypes: Okiep, Stein, Kleinsee and a Naries 1 individual (Fig. 3a). Okiep and Stein are found in the North of the distribution of clade B of *G. diffusa*, adjacent to the Roub morphotype, while Kleinsee is the only coastal morphotype, found along the Atlantic between the village of Kleinsee in the North and Hondeklip Bay in the South. Okiep and Kleinsee are very similar in terms of floral phenotype, but they differ strongly in their life histories since Kleinsee tends towards perenniality while Okiep is always annual, as are all the other morphotypes. Clade D, sister to clade C, contained the center of the diversity with eight morphotypes, seven occurring in a small geographic area from the town of Springbok in the North to Garies in the South and the widely distributed Nieuw morphotype occurring from the Cederberg region in the South to the Knervslakte in the North. Clade D comprised three well supported subclades; the first grouping Koma and Naries morphotypes, the second grouping the highly sexually deceptive morphotypes (Spring and Buffels), and the third grouping the southernmost morphotypes (Soeb, Garies, Cal and Nieuw). However, the relationship between these three subclades was not well supported. Koma and the Naries 2 individual are sister to each other, and together are inferred as sister to the rest of the morphotypes with weak support. In the high level of missingness topologies (90% to 70% missingness), Naries is therefore always polyphyletic, either clustering with Okiep or Koma. However, at lower levels of missingness, the two Naries individuals are found together in a clade with Koma (60% to 10% missingness except 40%) or Spring (40% missingness). Spring and Buffels are well supported as sister taxa, and together are weakly supported as sister to the grouping of southernmost morphotypes (Cal, Garies, Soeb and Nieuw). Garies appears polyphyletic with the two southernmost sampled sites (Garies 1 and 2) being inferred as sister to the Nieuw clade and the Garies 3 population inferred as sister to Cal. When using the complete dataset (*i.e*. with individuals close to hybrid zones), there were no significant changes in the general topology. One notable difference was that the morphotype Garies was even more polyphyletic since the Garies 4 population was nested into the Soeb clade. Also, in clade C, Kleinsee showed very short branch length between the different individuals sampled, further highlighting the high levels of relatedness between these individuals (see population structure section).

#### Coalescent-based approach

We were able to reconstruct the species tree using the Astral-III approach using between 4419 (90% missing data) and 491 (10% missing data) gene trees. The support values were generally high in the different topologies, confirming the results of the concatenation approach and further indicating that the different morphotypes formed distinct entities. As for the concatenation analysis, support decreased for species trees reconstructed with fewer loci. Apart from the lowest level of missing data (10%), all topologies were identical and mostly concordant with the concatenation approach (Fig. 3b). However, the main surprise comes from the placement of *G. corymbosa* which is no longer inferred as a sister species to *G. diffusa*, but is placed in a clade with the Northern morphotypes Rich, Khubus and Roub. As in the concatenation approach (Fig. 3a), the support for the relationships between the morphotypes Koma, Naries, Buffels and Spring found in clade D is low. However, the strongly sexually deceptive morphotypes Spring and Buffels are always inferred as sister morphotypes with strong support, as are the southernmost morphotypes (Cal, Garies, Nieuw, Soeb). The normalized quartet scores for each dataset were between 55% and 59%, indicating high levels of discordance in gene trees since only a little more than half of the gene trees supported the inferred species tree. Running Astral-III with the sampling location as grouping variable instead of morphotype, provided insight into the morphotypes inferred as polyphyletic in the concatenation based-approach (Naries and Garies). It showed that the two Naries populations found to fall in different clades in the concatenation analysis are inferred with very high support as a monophyletic clade. However, Garies remained polyphyletic and showed the same branching order as in the concatenation approach.

### Investigating causes of gene tree discordance

Of the 560 morphotype trios investigated for ABBA-BABA comparisons, 97 showed a *p*-value smaller than 0.05. However, after applying the Bonferroni-Holm correction, only 10 trios remained significant (Table 2). We classified these significant trios according to whether the groups involved in the trios were in the same *G. diffusa* geographic clade, in different *G. diffusa* clades or in different *Gorteria* species. 8 out of the 10 significant trios concerned introgression between clades of *Gorteria diffusa*. Among these, 6 of them involved the morphotype Roub (clade A) with introgression between this morphotype and Stein (clade C) and members of clade D (Soeb, Garies, Nieuw). This could explain the intermediate position of the Roub morphotype in the population genomic analyses. Only one significant trio involved *Gorteria corymbosa (i.e*. different species), and one involved Spring, Buffels and Garies, all found in clade D. Surprisingly, morphotypes known to hybridize in natural contact zones were not retrieved in this analysis.

**Table 2:**
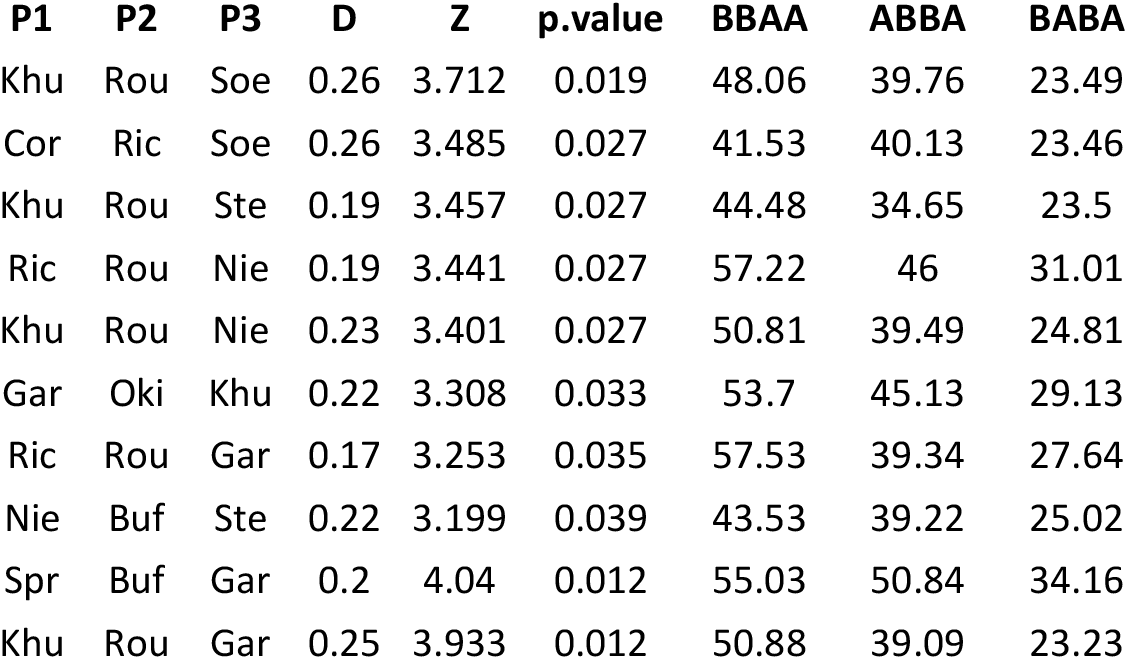
Results of the D-suite analysis. Each row represents a trio and its associated statistics.

| P1 | P2 | P3 | D | Z | p.value | BBAA | ABBA | BABA |
| --- | --- | --- | --- | --- | --- | --- | --- | --- |
| Khu | Rou | Soe | 0.26 | 3.712 | 0.019 | 48.06 | 39.76 | 23.49 |
| Cor | Ric | Soe | 0.26 | 3.485 | 0.027 | 41.53 | 40.13 | 23.46 |
| Khu | Rou | Ste | 0.19 | 3.457 | 0.027 | 44.48 | 34.65 | 23.5 |
| Ric | Rou | Nie | 0.19 | 3.441 | 0.027 | 57.22 | 46 | 31.01 |
| Khu | Rou | Nie | 0.23 | 3.401 | 0.027 | 50.81 | 39.49 | 24.81 |
| Gar | Oki | Khu | 0.22 | 3.308 | 0.033 | 53.7 | 45.13 | 29.13 |
| Ric | Rou | Gar | 0.17 | 3.253 | 0.035 | 57.53 | 39.34 | 27.64 |
| Nie | Buf | Ste | 0.22 | 3.199 | 0.039 | 43.53 | 39.22 | 25.02 |
| Spr | Buf | Gar | 0.2 | 4.04 | 0.012 | 55.03 | 50.84 | 34.16 |
| Khu | Rou | Gar | 0.25 | 3.933 | 0.012 | 50.88 | 39.09 | 23.23 |

### Ancestral trait reconstruction

The equal rates models were selected for all traits studied. We obtained good support for the traits thought to be responsible for variation in pollinator behavioural responses to *G. diffusa* morphotypes. First, the absence of a black spot on the ray floret was inferred as the ancestral character to the whole *Gorteria* clade. To further support this, *G. integrifolia* and *G. warmbadica*, the other known southern African *Gorteria* species (but not included in this study), which sit outside the *G. personata-G. corymbosa-G. diffusa* clade (Stångberg et al., 2013a), are also non-spotted species. The spotted character appeared first outside *G. diffusa* as *G. personata* and *G. piloselloides* have a full ring of spots. The presence of spots on only a fraction of the ray florets (“few spots”), an important innovation observed only in *G. diffusa*, was inferred as being the ancestral state of *G. diffusa*. Within *G. diffusa*, there was one secondary loss of the spotted character in the morphotype Khubus and the morphotypes with a full ring of spots were all found in clade D. Within clade D, the clade containing 3 out of the 4 ‘ringed’ morphotypes was inferred as presenting a full ring ancestrally, with a secondary loss of this character in the Nieuw morphotype. Second, yellow appeared as the ancestral color of *G. diffusa* flowers, with all other *Gorteria* species and clade A of *G. diffusa* being yellow-flowered, the orange colouration appeared as an innovation in *G. diffusa* and was inferred to be the ancestral state of clade B with two secondary changes to yellow (Stein and Soeb). Third, the presence of petal spot papillae, a character thought to contribute importantly to the sexual behaviour of *Megapalpus capensis*, appeared in *G. diffusa* in clade D and is inferred as the most likely ancestral state to this clade, with two secondary losses in Naries and Soeb.

Finally, the reconstruction of pollinator behaviour across the phylogeny indicates that sexual deception, as exemplified by pseudo-copulation of *M. capensis*, is a derived strategy that arose twice in the species (within clade D). Inspection behaviour (where male flies inspect the capitulum apparently searching for mates, but do not attempt to copulate) seems to have evolved also twice independently in two very different looking morphotypes: Cal and Okiep.

### Phylogenetic dating

The resulting chronogram from the BEAST2 analysis revealed a Pliocene-Quaternary diversification of the genus *Gorteria*, and an age for the most recent common ancestor of all *G. diffusa* morphotypes in the Quaternary (95% highest posterior density interval 632Kya - 2.25Mya) (Fig. 4).

**Figure 4:**
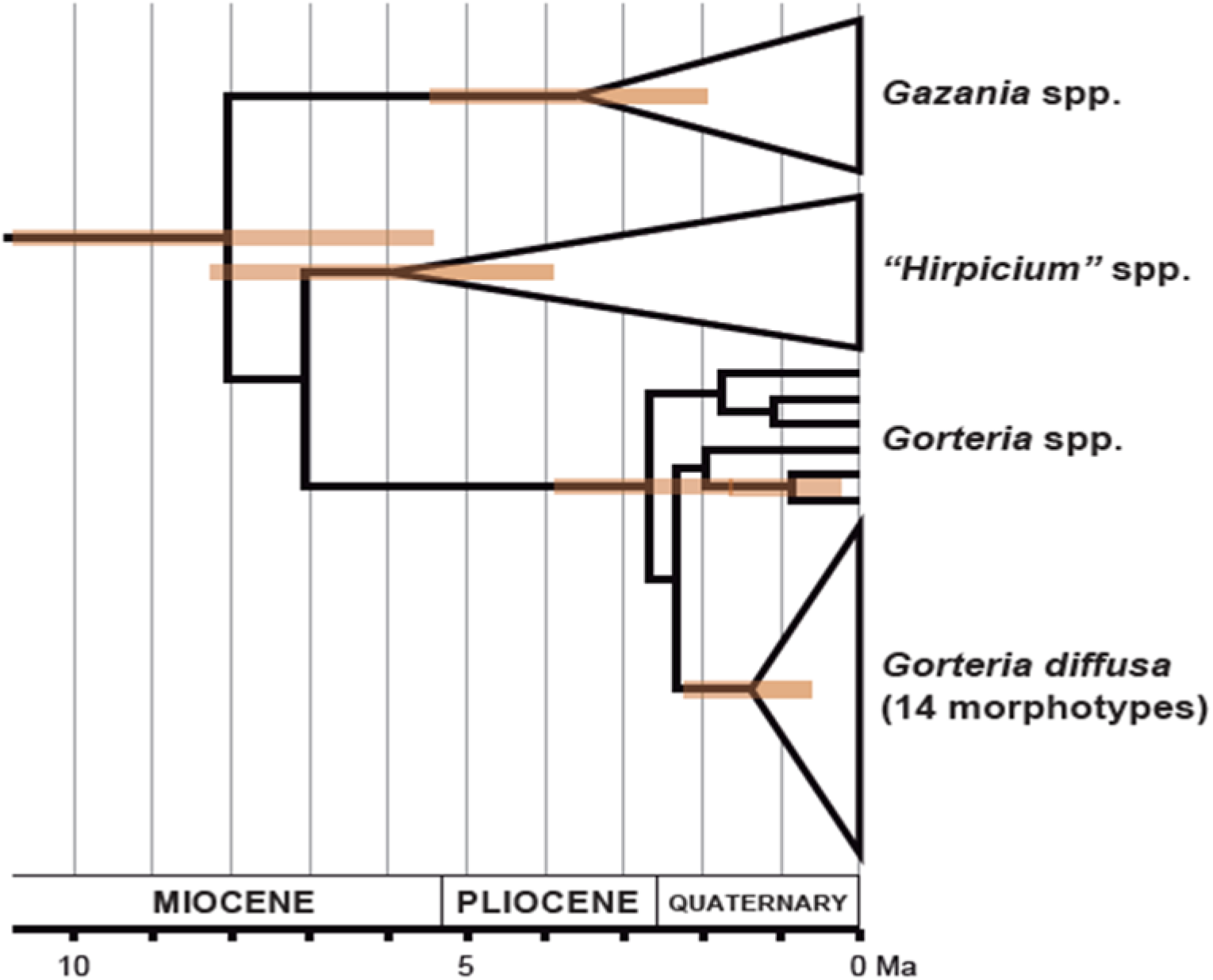
Results of the phylogenetic dating for the subfamily Cichorioideae using BEAST2. Coloured bars correspond to 95% of the posterior distributions of the age of each node with posterior probability > 0.95.

## Discussion

Using high-throughput sequencing techniques we were able to reconstruct the evolutionary history of this remarkable incipient radiation of *G. diffusa* morphotypes. Evidence from population genomic and phylogenomic analyses showed that the different morphotypes represent distinct genetic entities that have radiated recently. The detailed analysis of the different gene tree topologies highlighted the presence of high levels of incongruency likely due to the molecular technique used, incomplete lineage sorting and introgression between lineages. In addition, using ancestral trait reconstruction methods, we showed that sexual deception is likely a derived strategy that arose twice within the species complex, and which involves derived combinations of floral traits with different evolutionary histories.

### *Evolutionary history of the incipient radiation of* G. diffusa

Contrary to previous attempts to reconstruct the phylogenetic relationships within *G. diffusa* (Stångberg et al., 2013b), both methods used here (concatenation- and coalescent-based) allowed us to obtain a generally well-resolved tree both at the scale of the genus and at the intraspecific scale for *G. diffusa*. The main difference between the two approaches at the interspecific level concerned the position of *G. corymbosa*. In the concatenation-based approach, *G. diffusa* was confirmed as a monophyletic group with *G. corymbosa* as its sister species. In the coalescent-based analyses, *G. corymbosa* was found to be nested within the most basal Northern clade of *G. diffusa* (clade A in the concatenated analysis, Fig. 3). *G. corymbosa* inflorescences look morphologically similar to the Rich and Khubus morphotypes as they are the only *G. diffusa* forms with eight ray florets, sharing this characteristic with other outgroup species (*G. personata* and *G. piloselloides*). However, there are important characters distinguishing *G. diffusa* from *G. corymbosa*, most notably *G. corymbosa* always shows setiform phyllaries while phyllaries are triangular in *G. diffusa* (Stångberg and Anderberg, 2014). Leaves also tend to be different as they are always entire in *G. corymbosa* while *G. diffusa* tends to have a mix of entire and pinnatifid leaves (Stångberg et al., 2013a; Stångberg and Anderberg, 2014). Besides these morphological differences, previous phylogenetic work at the scale of the genus has supported *G. corymbosa* as a distinct species sister to *G. diffusa* (Stångberg et al., 2018, 2013a). This suggests that the placement of *G. corymbosa* within *G. diffusa* observed in our coalescent-based analysis could be spurious and due to external processes known to affect such methods. In particular, the *G. corymbosa* range abuts that of the Rich morphotype in the Ritchersveld region, suggesting that hybridization could happen between the two species. Hybridization could be responsible for the observed incongruence as gene flow violates the underlying assumptions of methods such as Astral-III and has been shown to generate inconsistent species trees (Solís-Lemus et al., 2016). Thus, while the bulk of available evidence supports *G. corymbosa* as sister to *G. diffusa*, the possibility that it is the northern-most morphotype of *G. diffusa* cannot be excluded at present. However, none of the significant introgression tests (Table 3) involved *G. corymbosa*. Further data, such as detailed morphological comparisons of the northern *G. diffusa* morphotypes and *G. corymbosa* or the addition of the *Gorteria* species not included in this study, would be required to resolve the placement of *G. corymbosa*.

At the within *G. diffusa* scale, both approaches allowed us to obtain a well-supported phylogenetic hypothesis with well-defined clades and most of the morphotypes being inferred as monophyletic. We retrieved the Northern–Southern structure (clade A and B, Fig. 3) suggested by previous phylogenetic work (Stångberg et al., 2013a). This basal split was not surprising considering that Rich and Khubus from the Northern clade are distinct from other *G. diffusa* morphotypes in having eight ray florets, triangular inner involucral bracts and globose mature involucres (Stångberg et al., 2013a). However, Roub, the third member of the Northern clade, has a different number of ray florets (mean of 13, as in all the morphotypes in clade B) and even if its placement within the Northern clade is supported in both phylogenetic approaches, its intermediate position in the population genomics analysis (Fig. 2) suggests that this could be a hybrid between clade A and clade B (Fig. 2, 3). The results of the ABBA-BABA analysis suggest that Roub is involved in introgression events with both members of clade A and clade B, further reinforcing this hypothesis. Geographically, Roub lies in between Rich (Clade A) and Stein (clade B) and further sampling along the geographic transition between these clades will be necessary to shed light on the origin of the Roub morphotype. Within clade B, clade C contained the only perennial form of *G. diffusa* (Kleinsee) but this did not stand out as particularly distinct from the rest of the morphotypes, and was always grouped with Okiep and Stein with good support (Fig. 3). The largest clade within *G. diffusa* (clade D) contained the center of the diversity with 8 morphotypes found in a relatively small geographic area (Fig. 1, Fig. 3). Within this clade, two morphotypes (Garies and Naries) were sometimes inferred as paraphyletic and the branching order of Koma, Naries and the Spring-Buffels clade was variable between analyses and showed smaller support values. Two main processes could explain these uncertainties. First, the morphotypes found in clade D are distributed in parapatry with many contact zones between them (Fig. 1). At these contact zones, individuals with intermediate phenotypes can be found, suggesting that there is some gene flow between the different morphotypes (Ellis and Johnson, 2009). However, ABBA-BABA analyses did not provide strong support for ubiquitous introgression amongst clade D morphotypes, even those known to currently form hybrids. Second, our phylogenetic dating analysis supported a recent diversification event within *G. diffusa* starting between 652kya and 2.25Mya, and therefore the likelihood of incomplete lineage sorting is high. These two processes, gene flow and incomplete lineage sorting, have been shown to generate inconsistencies in concatenation and coalescent-based approaches and discrepancies between them (*e.g*. (Kubatko and Degnan, 2007; Pollard et al., 2006; Roch and Steel, 2015; Solís-Lemus et al., 2016). Using more genomic markers, notably whole-genome assemblies, has been shown to be a powerful approach (see *e.g*. (Suvorov et al., 2021a; Vanderpool et al., 2020) and would enable us to resolve this part of the tree and better understand the processes that have led to this rapid diversification.

Overall, our phylogenetic results suggest the need for more detailed taxonomic work on the *G. diffusa* complex. We have shown that most morphotypes, defined on the basis of floral traits, are indeed well supported monophyletic entities, and that the complex comprises at least three well-defined clades that are geographically separated and each contain multiple floral morphotypes. In addition, the narrowness of the hybrid zones between morphotypes observed in the field and the placement of the hybridizing pairs in distinct clades suggests that reproductive isolation mechanisms are at play. Splitting the morphotypes into several species or subspecies would have important conservation implications as some of them have geographically restricted ranges and could be considered as being micro-endemic. Nevertheless, an important criterion to define distinct species would be to show that, when in contact, morphotypes still represent distinct genotypic clusters and do not fuse (Mallet, 2020, 1995). Using more sampling locations and performing detailed analyses of the contact zones, investigating patterns of genomic differentiation and mechanisms of reproductive isolation, will all be important steps towards this needed revision.

### Causes of gene tree incongruencies

We showed high levels of incongruencies between the different gene trees as only 55 to 59% of the gene trees supported the inferred species tree in the coalescent analysis. Genotyping-by-Sequencing generates short loci (ca. 150bp) which can generate errors in the reconstructed gene trees (Zhang et al., 2018). However, the molecular technique is unlikely to be the sole factor generating inconsistencies between gene trees. First, using a Bayesian phylogenetic dating method, we highlighted that the diversification of morphotypes took place recently, a factor strongly favouring incomplete lineage sorting as a likely explanation for gene tree incongruence (Degnan and Rosenberg, 2006). This is also likely the reason why previous attempts at reconstructing the history of diversification of the species were unsuccessful (Stångberg et al., 2013a). Second, several hybrid zones are found between pairs of parapatric morphotypes which is likely leading to effective introgression between morphotypes. Moreover, using ABBA-BABA tests, we found significant signals of introgression at various scales: between *Gorteria* species, between *G. diffusa* clades and within *G. diffusa* clades. While only 10 out of the 560 tests performed remained significant after correction for multiple testing, it is likely that this was the result of little statistical power due to the relatively low number of markers used and the high number of comparisons. *Gorteria diffusa* diversification was therefore likely influenced by various events of introgression. Interestingly, there is increasing evidence that introgression plays an important role in rapid diversification events such as adaptive radiation (*e.g*. (Malinsky et al., 2018a; Suvorov et al., 2021b), see also (Marques et al., 2019)). However, determining whether diversification in *G. diffusa* has been fueled by hybridization will require denser genomic data.

### Sexual deception

Obtaining a robust phylogenetic hypothesis at the within *G. diffusa* scale allowed us to map different phenotypic traits and infer the likely pattern of evolution of sexual deception. *Megapalpus capensis* behaviour in response to the different phenotypes is highly variable in the species, varying from simple feeding behaviour to strong copulation responses, with intermediate responses such as mate-searching without copulation (Ellis et al., 2014; Ellis and Johnson, 2010). The phenotypes eliciting the strongest sexual deception behaviour (*i.e*. copulation) all share the same combination of phenotypes, being orange with complex spots (*i.e*. with papillae) only on a few of their ray florets (Spring, Buffels, Nieuw and Koma) (Ellis et al., 2014). Interestingly, this combination of phenotypes appeared as phylogenetically derived in the *G. diffusa* radiation and our analyses suggested that this is the result of a combination of traits that have evolved sequentially in the radiation. First, the presence of anthocyanin spots on all ray florets first appeared outside the species and is present in some of the sister species (*e.g. G. piloselloides* and *G. personata*). It is also a common trait in the daisy communities found in Namaqualand, as it is found in various species of the *Arctotidae* tribe to which *Gorteria* belongs (*e.g*. in *Gazania* and *Arctotis*), and *Calendulae (e.g. Dimorphotheca*) and *Anthemidae (e.g. Ursinia*). This trait is likely of adaptive significance since it has been experimentally demonstrated that a ring of spots on ray florets significantly increased flower salience when found in a community with various phenotypes (de Jager et al., 2017), such as the large flower blooms observed in Namaqualand. Second, the presence of spots on a subset of the ray florets (usually 1 to 4) is restricted to *G. diffusa* and likely evolved in the common ancestor of the species (Fig. 5). A second origin of few spots is inferred for the Nieuw morphotype in the southern clade of *G. diffusa* following reversion to a ring of spots at the base of this clade. To the pollinator eye, the spots appear as scattered on the inflorescence, which could reinforce the function of the spot as a pollinator mimic. Flies have been shown to display an innate preference for dark spots related to their aggregating and mating behaviour (Eisikowitch, 1980; Goulson et al., 2009). This trait could have evolved using a rather simple genetic mechanism triggering the production of the spots only on the first ray florets to mature, rather than on all florets during development (Thomas et al., 2009). Third, the evolution of orange ray colour is inferred at the base of clade B, although it is also present in some populations of the Rich morphotype (Clade A) and *G. piloselloides*. *Megapalpus* flies have strong preference for orange daisies (Ellis et al., 2021) and indeed yellow morphotypes of *G. diffusa* such as Rich, Soeb and Stein are less exclusively associated with *Megapalpus* (Ellis and Johnson, 2009). Indeed, the dating of diversification of *G. diffusa* (632Kya - 2.25Mya) aligns well with the migration of *Megapalpus* into the Namaqualand landscape ca. 2.40 (1.09–5.67) million years ago (de Jager and Ellis, 2017). Finally, the evolution of the unique papillae cells within the anthocyanin spots appeared later in the *G. diffusa* radiation (clade D). This trait seems to be at the core of the sexual deceit behaviour elicited in the males of *Megapalpus capensis* as they strongly prefer the most complex spots (de Jager and Ellis, 2012).

**Figure 5:**
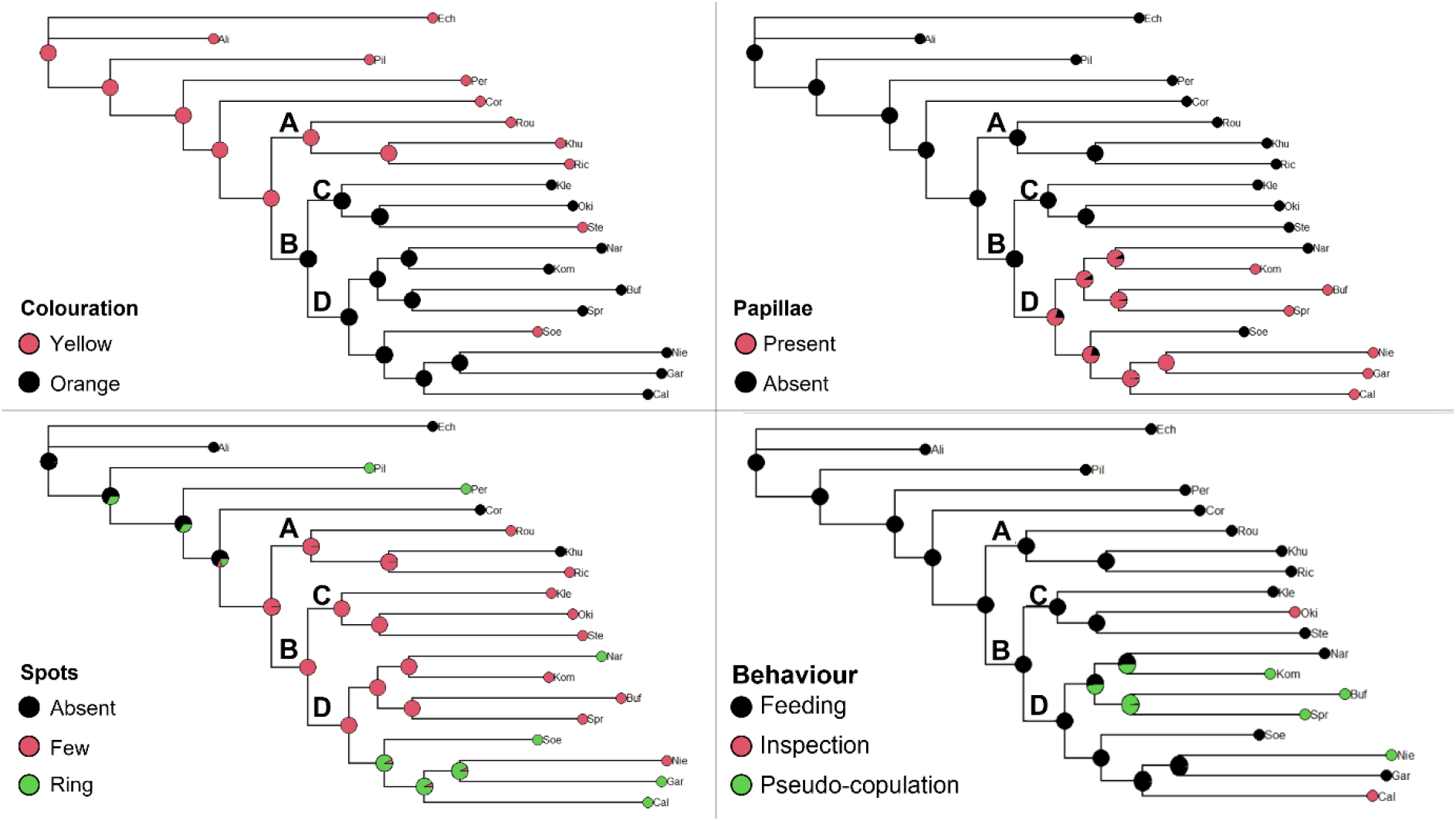
Results of the ancestral trait reconstruction. Colouration at each tip corresponds to the phenotype of each morphotype. At each node, the pie chart corresponds to the inferred probability of the ancestral phenotype.

The understanding of the genetic regulations of these different traits (anthocyanin spots, number of spots, colouration of the florets and presence of papillae) will enable us to understand how the different genetic pathways contributing to floral differentiation in the system are organized and how they became coupled together in certain morphotypes. Overall, there were no clear synapomorphies in the traits studied, involving phenotypic convergence, a pattern that is often explained by introgression (Stern, 2013), a mechanism which is highly likely in *G. diffusa* considering the known hybrid zones and results from the ABBA-BABA tests. Together with the detailed genetic regulation of the spot phenotypes, future whole genome assemblies of each morphotype will allow an understanding of which part of the genome have introgressed between the different morphotypes.

## Conclusion

In this study, we were able to resolve the history of diversification and trait evolution in one of the very few cases of sexual deceptive pollination outside of orchids. Our findings also highlight the power of reduced-representation techniques to solve recalcitrant phylogenetic problems such as recent evolutionary radiations. Future transcriptomic and genomic analyses performed on the different morphotypes will aid in understanding the genetic basis of the phenotypic differences as well as the genomic architecture of divergence and reproductive isolation.

## Acknowledgments

The authors thank Rachel Walker, Jurene Kemp, Caroli de Waal and Marinus de Jager for helpful discussions. We further thank the Northern Cape Department of Environment and Nature Conservation for issuing a collection permit. This study was supported by a Natural Environment Research Council (NERC) grant to B.J.G. and A.G.E. (NE/P011764/1), a Biotechnology and Biological Sciences Research Council (BBSRC) DTP studentship to G.M. and the Swiss National Science Foundation (mobility grant P2ZHP3_178043 to RTK).

## Supplementary Material

**Figure S1:**

**a.** Principal components analysis of the 824 SNPs shared by 80% of the *Gorteria diffusa* individuals. Outgroup species were not included in this analysis. Colour and shape correspond to the different morphotypes, and the name of the sampling localities are specified next to the position of each individual. First and third axis are represented, corresponding respectively to 9.6% and 4.4% of explained variance.

**b.** Principal components analysis of the 824 SNPs shared by 80% of the *Gorteria diffusa* individuals. Outgroup species were not included in this analysis. Colour and shape correspond to the different morphotypes, and the name of the sampling localities are specified next to the position of each individual. First and fourth axis are represented, corresponding respectively to 9.6% and 3.8% of explained variance.

**c.** fineRADstructure coancestry matrix. The colour of each square represents the results of a pairwise comparison of estimated coancestry between two individuals based on similarity between RAD loci.

The relative coancestry is illustrated, with high levels indicated by blue colouration and lower levels indicated by yellow. The tree is illustrative of relationships between populations but should not be interpreted as a phylogeny.

**Figure S2:**

**A)** Top panel: Boxplots representing the distribution of the ultra-fast bootstrap support values for each dataset. Dashed red line corresponds to the cut-off used for characterizing strong support.

Middle panel: Boxplots representing the distribution of the SH-aLRT support values for each dataset. Dashed red line corresponds to the cut-off used for characterizing strong support.
Bottom panel: Pie charts representing the number of supported nodes (SH-aLRT > 80% & UFB > 95%, light grey) against non-supported ones (dark grey).

**B)** Multidimensional scaling representation of the pairwise topological distance between the different assemblies at the 90% missing data threshold.

**Table S1:** Number of retained reads per individual.

| Sample | Number of retained reads |
| --- | --- |
| Ric_2B | 9065220 |
| Cal_4A | 7018967 |
| Cal_3A | 6964861 |
| Cal_2B | 6742335 |
| Gar_2A | 6678846 |
| Ste_2A | 6655208 |
| Cal_1C | 6155151 |
| Cal_3B | 5993641 |
| Ric_1C | 5905811 |
| Spr_1A | 5733324 |
| Oki_1B | 5604986 |
| Gar_1B | 5277001 |
| Cal_2A | 5150741 |
| Kom_3A | 5126454 |
| Ric_1B | 4949710 |
| Nie_4B | 4663211 |
| Khu_1A | 4600930 |
| Hir_1B | 4358077 |
| Kom_1C | 4272700 |
| Soe_2A | 3931451 |
| Spr_4B | 3895440 |
| Khu_2C | 3827133 |
| Soe_2C | 3797881 |
| Kom_3B | 3775695 |
| Kom_2C | 3774006 |
| Cal_1A | 3621136 |
| Rou_6A | 3563517 |
| Oki_3A | 3450602 |
| Spr_3B | 3449278 |
| Nie_3B | 3404405 |
| Buf_2B | 3320678 |
| Wor_3B | 3273382 |
| Ste_2B | 3176742 |
| Cal_4B | 3168449 |
| Nie_4C | 3137916 |
| Kom_1B | 3129817 |
| Rou_5C | 3059481 |
| Soe_1A | 3049346 |
| Spr_2B | 3032267 |
| Gar_1A | 2935134 |
| Buf_1B | 2923431 |
| Oki_2A | 2883564 |
| Nar_2C | 2813291 |
| Oki_1A | 2756335 |
| Kom_2A | 2675213 |
| Nie_2C | 2649743 |
| Per_2A | 2585066 |
| Buf_1A | 2468054 |
| Soe_1C | 2393623 |
| Oki_2B | 2323460 |
| Wor_2A | 2323164 |
| Gar_3C | 2277455 |
| Oub_2A | 2213105 |
| Cor_1B | 2205267 |
| Khu_2B | 2189654 |
| Soe_4A | 2166229 |
| Ric_3B | 2030977 |
| Nie_1A | 2007519 |
| Rou_6C | 1950769 |
| Gar_3A | 1415893 |
| Kle_1B | 1270467 |
| Cor_1A | 1208464 |
| Spr_4A | 1087080 |
| Wor_1C | 1080269 |
| Per_1C | 941663 |
| Hec_1A | 869772 |
| Ric_3A | 811628 |
| Spr_2A | 794634 |
| Spr_3C | 679798 |
| Kle_1C | 650050 |
| Kle_1A | 648168 |
| Gar_4C | 643892 |
| Per_2C | 563948 |
| Nar_1B | 523796 |
| Buf_3C | 501847 |
| Oki_3B | 467752 |
| Spr_1C | 384151 |
| Soe_3B | 357267 |
| Nar_3B | 313164 |
| Gar_4B | 311086 |
| Nar_2A | 181216 |
| Nar_1A | 143917 |
| Spr_2C | 129536 |
| Oub_3C | 106588 |
| Nar_3C | 34183 |
| Wor_2B | 24012 |
| Ste_1C | 18726 |
| Buf_2A | 17722 |
| Ste_1B | 10989 |
| Soe_4C | 9893 |
| Soe_3C | 4964 |
| Oub_4A | 2857 |

**Table S2.** Species of Cichorioideae (Asteraceae) included in the phylogenetic dating analysis and GenBank accession numbers of ITS and *trnL*-*trnF* sequences. Newly generated sequences are shown in bold.

| **Species** | **ITS** | ***trnL-trnF*** |
| --- | --- | --- |
| *Albertinia brasiliensis* | EF155744 | EF155832 |
| *Andryala integrifolia* | AY879153 | AY879119 |
| *Arctotheca calendula* | DQ889629 | DQ889645 |
| *Arctotis angustifolia* | EU846346 | EU846485 |
| *Arctotis lanceolata* | EU846370 | EU846509 |
| *Baccharoides adoensis* | EF155745 | EF155833 |
| *Baccharoides lasiopus* | EF155796 | EF155884 |
| *Berkheya angolensis* | EU527205 | EU527255 |
| *Berkheya annectens* | EU527211 | EU527261 |
| *Berkheya bipinnatifida* | EU527198 | EU527248 |
| *Berkheya canescens* | EU527201 | EU527251 |
| *Berkheya cardopatifolia* | EU527213 | EU527263 |
| *Berkheya carlinopsis* | AY504709 | AY504791 |
| *Berkheya cirsifolia* | EU527207 | EU527257 |
| *Berkheya coddii* | EU527194 | EU527244 |
| *Berkheya cruciata* | AY504712 | AY504794 |
| *Berkheya echinacea* | EU527204 | EU527254 |
| *Berkheya eriobasis* | EU527214 | EU527264 |
| *Berkheya fruticosa* | EU527199 | EU527249 |
| *Berkheya onobromoides* | EU527212 | EU527262 |
| *Berkheya pannosa* | EU527209 | EU527259 |
| *Berkheya pinnatifida* | EU527196 | EU527246 |
| *Berkheya rhapontica* | EU527206 | EU527256 |
| *Berkheya rigida* | EU527195 | EU527245 |
| *Berkheya setifera* | EU527208 | EU527258 |
| *Berkheya spinosa* | EU527202 | EU527252 |
| *Berkheya spinosissima* | EU527200 | EU527250 |
| *Berkheya subulata* | EU527203 | EU527253 |
| *Berkheya zeyheri* | EU527193 | EU527243 |
| *Bothriocline laxa* | EF155746 | EF155834 |
| *Brachythrix glomerata* | JN715915 | JN715872 |
| *Cabobanthus polysphaerus* | EF155749 | EF155837 |
| *Cacosmia harlingii* | JN837120 | JN837210 |
| *Centrapalus pauciflorus* | EF155751 | EF155839 |
| *Centratherum punctatum* | EF155753 | EF155841 |
| *Chionopappus benthamii* | JN837175 | JN837265 |
| *Chresta sphaerocephala* | EF155755 | EF155843 |
| *Chrysactinium acaule* | JN837158 | JN837248 |
| *Chrysolaena flexuosa* | EF155756 | EF155844 |
| *Cicerbita alpina* | LT721978 | KF486058 |
| *Cichorium intybus* | AJ633450 | GU817987 |
| *Crepidiastrum lanceolatum* | AB598563 | AB598612 |
| *Crepis aurea* | AF528483 | AF528396 |
| *Critoniopsis huairacajana* | EF155821 | EF155909 |
| *Cullumia aculeata* | EU527219 | EU527269 |
| *Cullumia bisulca* | EU527218 | EU527268 |
| *Cullumia decurrens* | EU527216 | EU527266 |
| *Cullumia patula* | EU527215 | EU527265 |
| *Cullumia rigida* | AY504714 | AY504796 |
| *Cuspidia cernua* | EU527220 | EU527270 |
| *Cymbonotus lawsonianus* | EU846333 | EU846473 |
| *Didelta carnosa* | EU527222 | EU527272 |
| *Didelta spinosa* | AY504717 | AY504799 |
| *Dillandia perfoliata* | JN837125 | JN837215 |
| *Distephanus polygalifolius* | HQ158399 | HQ158451 |
| *Dymondia margaretae* | AY504707 | AY504789 |
| *Eirmocephala brachiata* | EF155763 | EF155851 |
| *Elephantopus scaber* | JN715907 | JN715864 |
| *Erato polymnioides* | JN837164 | JN837254 |
| *Eremanthus erythropappus* | EF155770 | EF155858 |
| *Eremosis shannoni* | EF155758 | EF155846 |
| *Eremothamnus marlothianus* | JN837107 | JN837197 |
| *Ethulia conyzoides* | EF155772 | EF155860 |
| *Ferreyranthus verbascifolius* | JN837122 | JN837212 |
| *Gazania heterochaeta* | EU527228 | EU527278 |
| *Gazania krebsiana* | EU527225 | EU527275 |
| *Gazania lichtensteinii* | EU527226 | EU527276 |
| *Gazania longiscapa* | EU527227 | EU527277 |
| *Gazania rigens* | EU527229 | EU527279 |
| *Gazania rigida* | EU527230 | EU527280 |
| *Gazania tenuifolia* | AY504720 | AY504802 |
| *Gorceixia decurrens* | EF155773 | EF155861 |
| *Gorteria alienata* | JQ220237 | JQ220175 |
| *Gorteria corymbosa* | JQ220243 | JQ220183 |
| *Gorteria diffusa* Buffel | JQ220283 | JQ220219 |
| *Gorteria diffusa* Cal | JQ220294 | JQ220230 |
| *Gorteria diffusa* Garies | JQ220263 | JQ220199 |
| *Gorteria diffusa* Khubus | JQ220280 | JQ220216 |
| *Gorteria diffusa* Koma | JQ220259 | JQ220195 |
| *Gorteria diffusa* KZ | JQ220255 | JQ220189 |
| *Gorteria diffusa* Naries | JQ220292 | JQ220228 |
| *Gorteria diffusa* Nieuw | JQ220258 | JQ220194 |
| *Gorteria diffusa* Okiep | JQ220281 | JQ220217 |
| *Gorteria diffusa* Oubees | JQ220287 | JQ220223 |
| *Gorteria diffusa* Rich | JQ220269 | JQ220204 |
| *Gorteria diffusa* Soeb | JQ220260 | JQ220196 |
| *Gorteria diffusa* Spring | **OP422351** | **OP415050** |
| *Gorteria diffusa* Stein | **OP422352** | **OP415051** |
| *Gorteria integrifolia* | JQ220254 | JQ220191 |
| *Gorteria parviligulata* | JQ220264 | JQ220200 |
| *Gorteria personata* | JQ220267 | JQ220203 |
| *Gorteria piloselloides* | JQ220272 | JQ220208 |
| *Gundelia tournefortii* | AY504691 | AY504773 |
| *Gymnanthemum corymbosum* | JN715898 | JN715855 |
| *Haplocarpha lanata* | DQ889641 | DQ889657 |
| *Haplocarpha scaposa* | DQ889642 | DQ889658 |
| *Hesperomannia arbuscula* | JN837112 | JN837202 |
| *Heterocoma lanuginosa* | EF155798 | EF155886 |
| *Heterocypsela andersonii* | EF155779 | EF155867 |
| *Heterolepis aliena* | AY504700 | AY504782 |
| *Heterorhachis aculeata* | EU527235 | EU527285 |
| *Hieracium murorum* | AF528492 | AF528400 |
| *Hilliardiella leopoldii* | EF155781 | EF155869 |
| *Hilliardiella oligocephala* | EF155782 | EF155870 |
| *Hirpicium armerioides* | JQ220239 | JQ220176 |
| *Hirpicium bechuanense* | JQ220241 | JQ220177 |
| *Hirpicium echinus* | JQ220248 | JQ220184 |
| *Hirpicium gazanioides* | AY504726 | AY504808 |
| *Hirpicium gorterioides* | EU527240 | EU527290 |
| *Hirpicium linearifolium* | JQ220256 | JQ220192 |
| *Hirpicium pechuelii* | EU527239 | EU527289 |
| *Hoplophyllum spinosum* | JN837106 | JN837196 |
| *Hyoseris radiata* | AF528494 | AF528401 |
| *Hypochaeris radicata* | AF528461 | AF528381 |
| *Lactuca serriola* | KF485662 | KF486175 |
| *Launaea sarmentosa* | KF485537 | KF486048 |
| *Leiboldia guerreroana* | EF155820 | EF155908 |
| *Leontodon hispidus* | AF528485 | AF528393 |
| *Lepidaploa balansae* | EF155784 | EF155872 |
| *Lepidaploa borinquensis* | EF155785 | EF155873 |
| *Lessingianthus bardanoides* | JN715908 | JN715865 |
| *Liabum acuminatum* | JN837131 | JN837221 |
| *Linzia glabra* | JN715899 | JN715856 |
| *Microliabum polymnioides* | JN837168 | JN837258 |
| *Moquinia racemosa* | JN837117 | JN837207 |
| *Munnozia campii* | JN837152 | JN837242 |
| *Munnozia gigantea* | AY504697 | AY504779 |
| *Munnozia lyrata* | JN837157 | JN837247 |
| *Munnozia wilburii* | JN837153 | JN837243 |
| *Nabalus tatarinowii* | HQ436195 | HQ436129 |
| *Notoseris psilolepis* | KF485599 | KF486110 |
| *Oligactis sessiliflora* | JN837124 | JN837214 |
| *Orbivestus cinerascens* | EF155794 | EF155882 |
| *Paranephelium uniflorus* | AB355508 | AB355581 |
| *Parapolydora fastigiata* | EF155797 | EF155885 |
| *Philoglossa peruviana* | JN837163 | JN837253 |
| *Picris hieracioides* | KF154362 | KF196065 |
| *Platycarphella carlinoides* | AY504701 | AY504783 |
| *Prenanthes purpurea* | KF485548 | KF486059 |
| *Pseudonoseris szyszyłowiczii* | JN837172 | JN837262 |
| *Pseudostiffia kingii* | JN837116 | JN837206 |
| *Rhagadiolus edulis* | AF528495 | AF528402 |
| *Sampera cuatrecasasii* | JN837126 | JN837216 |
| *Scolymus maculatus* | AJ633469 | EU385110 |
| *Sinclairia discolor* | JN837181 | JN837271 |
| *Sinclairiopsis klattii* | JN837194 | JN837284 |
| *Sonchus oleraceus* | KF154364 | KF196067 |
| *Stokesia laevis* | EF155799 | EF155887 |
| *Stramentopappus pooleae* | EF155801 | EF155889 |
| *Strobocalyx arborea* | EF155774 | EF155862 |
| *Synclathium disciforme* | HQ436200 | HQ436134 |
| *Taraxacum officinale* | MG519289 | EF015611 |
| *Tarlmounia elliptica* | HQ158413 | HQ158465 |
| *Tragopogon pratensis* | KT167089 | KT167134 |
| *Vernonanthura chamaedrys* | JN715906 | JN715863 |
| *Vernonia angustifolia* | EF155807 | EF155895 |
| *Vernonia baldwinii* | EF155808 | EF155896 |
| *Vernonia brachycalyx* | EF155809 | EF155897 |
| *Vernonia bullata* | EF155810 | EF155898 |
| *Vernonia fasciculata* | EF155815 | EF155903 |
| *Vernonia profuga* | EF155826 | EF155914 |
| *Vernonia subplumosa* | EF155830 | EF155918 |
| *Vernonia texana* | EF155831 | EF155919 |
| *Vernoniastrum nestor* | EF155804 | EF155892 |
| *Warionia saharae* | AY190608 | AH013974 |
| *Youngia heterophylla* | KF154372 | KF196075 |

## Supplementary Material

### DNA extraction protocol

Silica dried leaf material was ground to a powder using glass beads and a Tissue Lyser II (Qiagen). The resulting powdered material was incubated at 55°C with CTAB buffer (Appendix One) before two chloroform washing steps and precipitation with sodium acetate and isopropanol. The resulting DNA was quantitatively and qualitatively analysed using a Qubit 2.0 Fluorometer (ThermoFisher) and 0.8% agarose gel electrophoresis, respectively.

### GBS library preparation

Genotyping-by-Sequencing library preparation followed approximately that of Escudero et al. (2014). 500ng of DNA per individual was digested with the restriction enzyme PstI (C_|_TGCA^v^G) for two hours at 37°C before quenching at 80°C for twenty minutes. Barcoded adapters (4-9bp) and common adapters (Appendix Two) were ligated to the resulting fragments (16°C, 2h) with the barcode sequences designed to reduce sequencing bias and favour nucleotide diversity (Herten et al., 2015). Samples were pooled and purified with Agencourt AMPure XP SPRI beads (Beckman-Coulter) followed by a low-cycle PCR amplification step (*Taq*, NEB 68°C 5min; 95°C 60s; 18x 95°C 30s, 65°C 30s and 68°C 30s; 68°C 5min) to reduce the levels of PCR duplicates in the library. The final library preparation was analysed using an Agilent 2100 Bioanalyser and the quantification validated using a Qubit 2.0 (ThermoFisher). High-throughput sequencing was performed on a single lane of 100bp paired-end sequencing on an Illumina HiSeq 2000 by the Beijing Genomics Institute (Copenhagen, Denmark).

