## Supplementary figures and images for "The phylogenetic history of the *Gorteria diffusa* radiation sheds light on the origins of plant sexual deception"

### Supplemental Figures

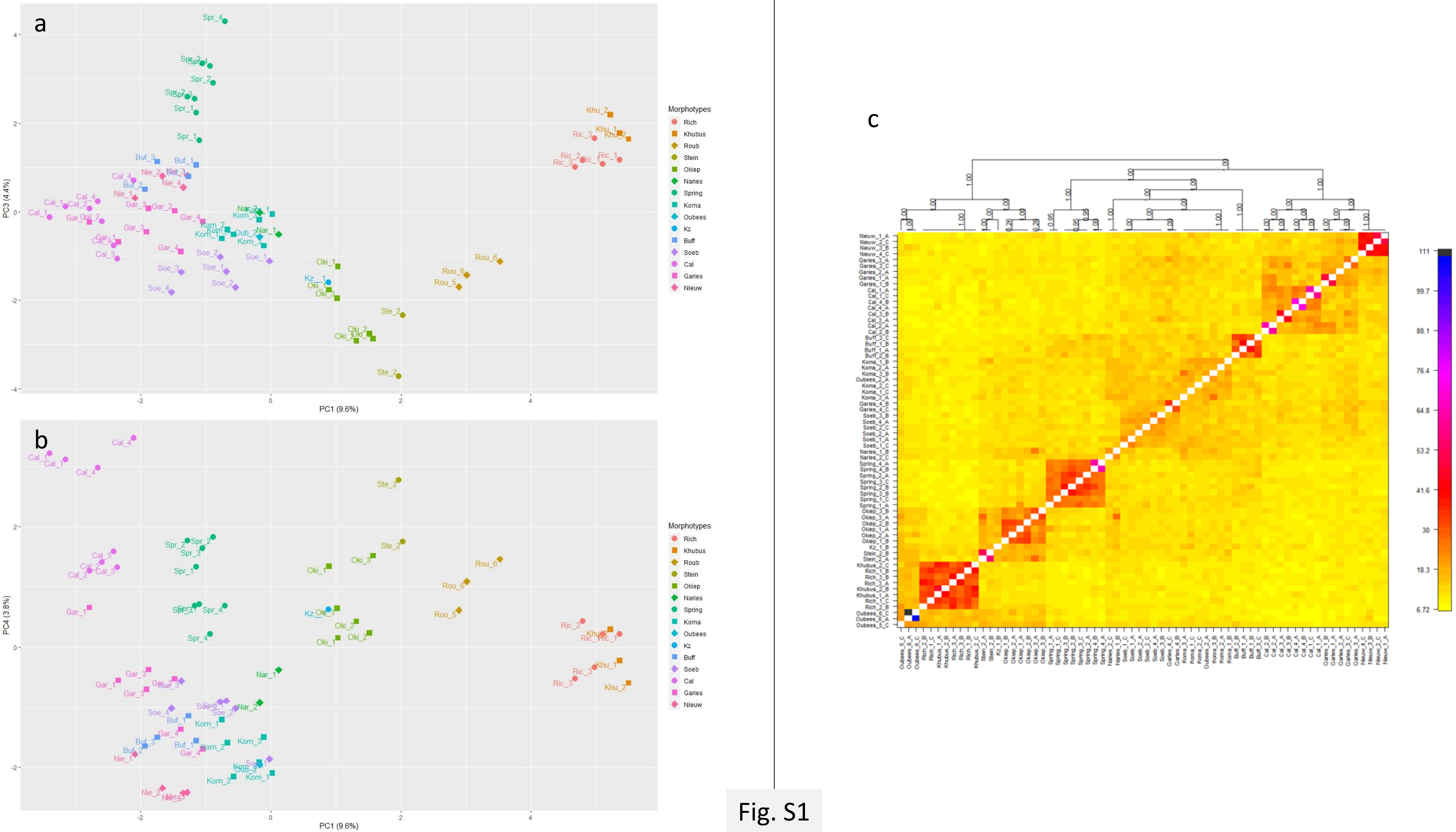

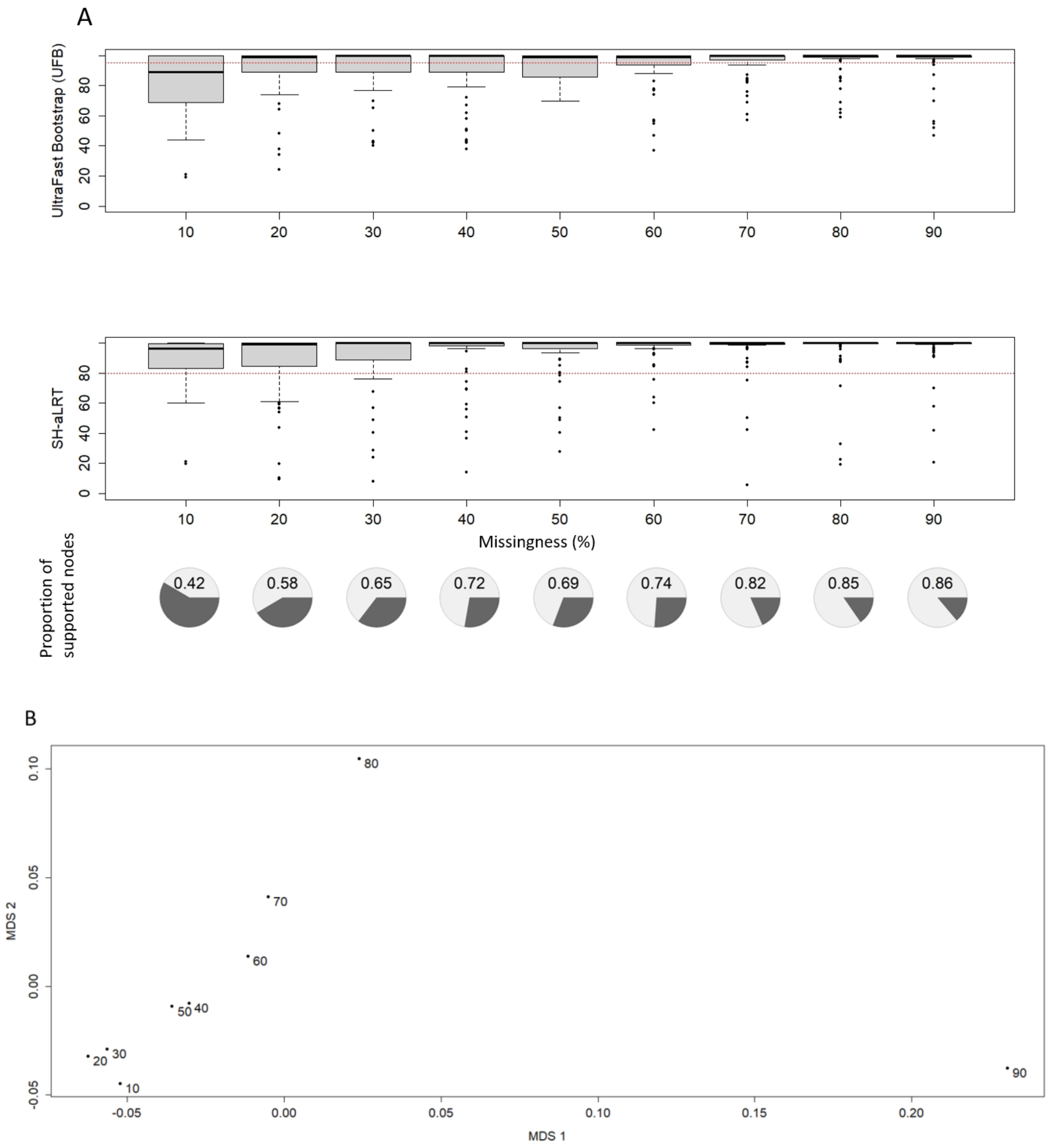
